# Voluntary motor commands are preferentially released during restricted sensorimotor beta rhythm phases

**DOI:** 10.1101/776393

**Authors:** Sara J Hussain, Mary K Vollmer, Iñaki Iturrate, Romain Quentin

## Abstract

Voluntary movement requires motor commands to be released from motor cortex (M1) and transmitted to spinal motoneurons and effector muscles. M1 activity oscillates between brief excitatory and inhibitory states that correlate with single neuron spiking rates. Here, we asked if the motor commands needed to produce voluntary, self-paced finger movements are preferentially released from M1 during restricted phases of this ongoing sensorimotor oscillatory activity. 21 healthy adults performed a self-paced finger movement task while EEG and EMG signals were recorded. For each finger movement, we identified the individual sensorimotor mu (8-12 Hz) and beta (13-35 Hz) oscillatory phase at the estimated time of motor command release from M1 by subtracting individually-defined MEP latencies from EMG-determined movement onset times. We report that motor commands were preferentially released at ~120° along the beta cycle but were released uniformly across the mu cycle. These results suggest that motor commands are preferentially released from M1 near optimal peak phases of endogenous beta rhythms.

## INTRODUCTION

Voluntary movement allows us to effectively interact with our environment and is central to human behavior. Such movements are produced when the primary motor cortex (M1) issues descending motor commands that are transmitted to spinal motoneurons and connected effector muscles. However, M1 activity is not static over time but oscillates in the mu (8-12 Hz) and beta (13-35 Hz) ranges (Pineda 2005; Pfurtscheller and da Silva 1999). These oscillations reflect rapidly alternating periods of excitation and inhibition (Buszaki 2006; Murthy and Fetz 1992; Murthy and Fetz 1996a; Murthy and Fetz 1996b; Salmelin and Hari 1994; Pfurtscheller and Aranibar 1979; Pfurtscheller and Lopes da Silva 1999) that correlate with local neuronal firing rates (Haegens et al. 2011; Murthy and Fetz 1996a), population-level neuronal activity (Miller et al. 2012) and communication between M1 and spinal motoneurons (Mima and Hallett 1999; Khademi et al. 2018; Khademi et al. 2019; Zrenner et al. 2018; Hussain et al. 2019; Bergmann et al. 2019; Torrecillos et al. 2020). Outside of the motor system, oscillatory phase shapes sensory and cognitive function, including perception (VanRullen 2016; Busch et al. 2009; Dugué et al. 2009; Hanslmayr et al. 2013; Baumgarten et al. 2015), attention (VanRullen 2018; Busch and VanRullen 2010) and memory (Kerrén et al. 2018; Ter Wal et al. 2020; Ten Oever et al. 2020). Yet, the relationship between sensorimotor oscillatory phase and voluntary motor behavior has remained largely unexplored.

Within M1, single-neuron spiking rates, population-level neuronal activity, and corticospinal motor output are all increased during trough phases of mu and beta rhythms (Murthy and Fetz 1996a; Haegens et al. 2011; Miller et al. 2012; Zrenner et al. 2018; Hussain et al. 2019; Bergmann et al. 2019). Based on these findings, it has been proposed that oscillations in the membrane potential of the layer V pyramidal neurons that causally produce voluntary movement (Brecht et al. 2004) generate phase-dependent fluctuations in M1 activity and its output (Zrenner et al. 2018; Hussain et al. 2019; Bergmann et al. 2019). Because motor commands are only released from M1 when its activity reaches an excitatory threshold (Hanes and Schall 1995; Chen et al. 1998), it is possible that phase-dependent fluctuations in M1 activity determine when along the mu and beta oscillatory cycle motor commands are most likely to be released. Here, we tested the hypothesis that motor commands are preferentially released from M1 during restricted phase ranges of sensorimotor mu and beta rhythms. We report that motor commands were preferentially released from M1 at ~120° along the beta cycle but were released uniformly across the mu cycle. Results demonstrate that motor commands are most often released from M1 within restricted, optimal beta phase ranges in the healthy human brain.

## Methods

### Data acquisition

#### Subjects and experimental design

21 healthy subjects participated in this study (11 F, 10 M, age = 27.66 ± 5.01 [SD] years). This study was approved by the National Institutes of Health Combined Neuroscience Section Institutional Review Board. Prior to participation, all subjects provided their written informed consent.

#### EEG and EMG recording

64-channel EEG signals (ground: O10; reference: AFz) and bipolar EMG signals (ground: dorsum of left wrist) were recorded using TMS-compatible amplifiers (NeurOne Tesla, Bittium, Finland) at 5 kHz (low-pass hardware filtering cutoff frequency: 1250 Hz, resolution: 0.001 µV) during single-pulse TMS delivery and the self-paced finger movement task. EEG impedances were maintained below 10 kΩ and EMG was recorded from the left first dorsal interosseous muscle (L. FDI) using disposable adhesive electrodes arranged in a belly-tendon montage.

#### Single-pulse TMS delivery

The scalp hotspot for the L. FDI muscle was identified over the hand representation of the right M1 as the site that elicited the largest MEP and a visible, focal twitch in the L. FDI muscle following suprathreshold single-pulse TMS. Resting motor threshold (RMT) was determined using an automatic threshold-tracking algorithm (Adaptive PEST Procedure; Awiszus and Borckhardt 2011). Then, 50 single TMS pulses were delivered to the L. FDI hotspot at 120% RMT (inter-pulse interval: 6 s with 0.9 s jitter). The MEP latency was defined as the amount of time needed for action potentials produced by TMS to travel from the stimulated M1 to the L. FDI muscle (see *EMG processing*). All TMS procedures were performed using a figure-of-eight coil held at ~45° relative to the mid-sagittal line (MagStim Rapid^2^, biphasic pulse shape; MagStim Co Ltd., UK). RMT was 61.62 ± 12.27% of maximum stimulator output.

#### Self-paced finger movement task

Subjects performed a self-paced voluntary finger movement task during which they viewed a series of unique pictures on a computer screen (Places Scene Recognition Database; Zhou et al. 2017, see Figure 1a). Subjects were instructed to view each picture for as long as they desired and then press a button on a standard keyboard (left CTRL button) using their left index finger when they wished to view the next picture in the series. Between pictures, a fixation cross was presented (inter-trial interval: 1.5 s with 0.2 s jitter). The task was designed to ensure that subjects produced a self-paced, discrete finger movement whenever they desired. During the task, the left arm was supported on a pillow to ensure full muscle relaxation and prevent extraneous movement. The finger movement task was divided into 6 blocks of 100 unique pictures (i.e., 6 blocks of 100 movements). Subjects were given a short break after 3 blocks. To ensure that subjects were not merely reacting as fast as possible to the presentation of each picture and that movements were truly self-paced, trials with reaction times faster than 400 ms were excluded from analysis.

**Figure 1.**
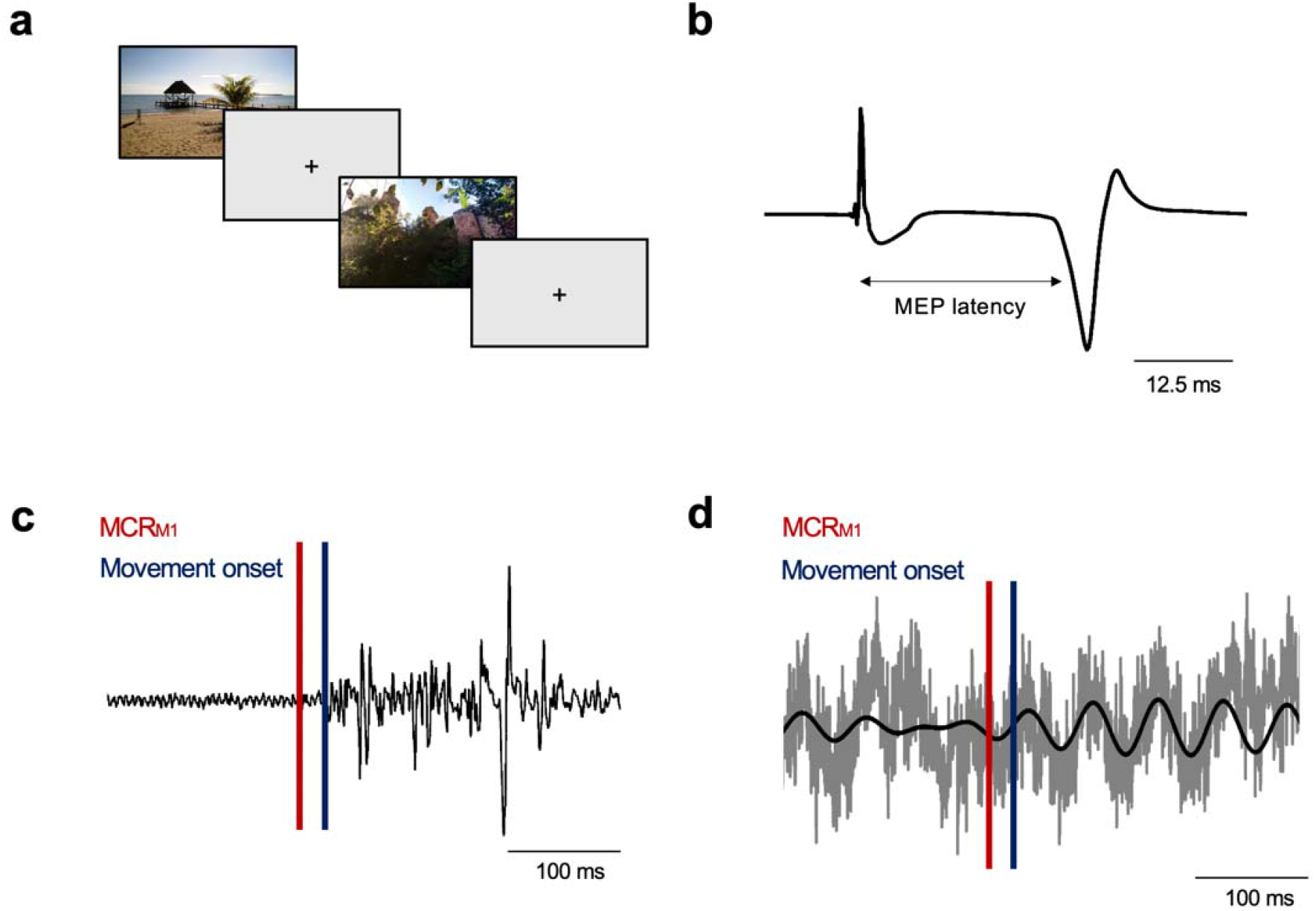
Experimental approach. a) Subjects viewed a series of pictures on a computer screen while they performed a self-paced finger movement task. Subjects were instructed to press a button using their left index finger whenever they wished to move to the next picture. EMG and EEG signals were recorded during the task. b) Single-pulse TMS was used to determine each subject’s individual left first dorsal interosseous (L. FDI) motor-evoked potential (MEP) latency. The large positive spike reflects the TMS pulse artifact. c) The time of motor command release from M1 (MCR_M1_ red vertical line) for each movement was estimated by subtracting each subject’s MEP latency from EMG-defined movement onset times (blue vertical line). The black trace reflects raw EMG data. d) Mu and beta oscillatory phase at the time of MCR_M1_ was then determined (see *Methods* for details). The grey and black traces reflect raw and filtered EEG data, respectively.

### Data analysis

#### EMG processing

EMG signals were processed using FieldTrip (Oostenveld et al. 2011) combined with custom-written scripts (MATLAB; TheMathWorks, Natick MA). EMG signals were used to: 1) measure individual MEP latencies obtained using single-pulse TMS, and 2) determine movement onset during the finger movement task.

To measure MEP latencies, data were divided into segments (−0.25 to -0.100 s relative to TMS pulse), demeaned, and linearly detrended. For each subject, MEP signals were averaged over trials to generate a mean MEP signal. The inflection point between post-stimulus baseline EMG activity and the beginning of the averaged waveform was visually identified as the MEP latency (see Figure 1b). For two subjects in whom MEP latencies could not be reliably identified, MEP latencies were set equal to the mean latency obtained in all other subjects. Average MEP latencies across subjects were 23.5 ± 1.64 ms.

To measure movement onset, EMG data were divided into segments (± 1.2 s relative to the button press during the task), demeaned, and linearly detrended. A notch filter (zero-phase shift Butterworth filter; order=4, 58-60 Hz) was applied to remove line noise, followed by a high-pass filter (zero-phase shift Butterworth filter; order=6, cutoff frequency: 20 Hz, De Luca et al. 2010). Voluntary EMG activity onset for each button press was visually identified using a combination of frequency- and time-domain EMG analysis, as previous studies have shown that visual inspection provides the most reliable results compared to automatic algorithms (Tenan et al. 2017). EMG signals were spectrally decomposed over time (multi-taper method using Hanning tapers, 20-500 Hz, time window=0.100 s, 0.2 ms resolution). To identify voluntary EMG onsets preceding each button press, time-frequency representations were plotted alongside EMG but not EEG signals. The experimenter identifying EMG onsets was thus blinded to EEG activity during EMG onset identification. Movements in which EMG onsets could not be reliably identified due to inadequate relaxation of the L. FDI between movements were excluded from analysis.

#### EEG processing

EEG signals were processed with FieldTrip (Oostenveld et al. 2011) and custom-written MATLAB scripts (TheMathWorks, Natick MA). Signals recorded during the finger movement task were divided into 2.4 s segments (± 1.2 s relative to the button press). Segmented data were re-referenced to the average reference, demeaned, and linearly detrended. Channels with impedances >10 kΩ were removed, and independent components analysis (ICA) was used to attenuate EEG artifacts. After ICA, channels previously removed due to high impedances were interpolated using spherical splines, and finally a subset of channels overlying the right sensorimotor cortex contralateral to the L. FDI (FC2, FC4, FC6, Cz, C2, C4, C6, T8, CP2, CP4, CP6) were selected for further analysis. To attenuate the effects of volume conduction, each selected channel and its four neighbors were used to obtain Hjorth-transformed signals (Hjorth 1975; Zrenner et al. 2018; Hussain et al. 2019).

We individualized the scalp channels used for mu and beta analysis by identifying those that best captured movement-related mu and beta reactivity (Torrecillos et al. 2020). using time-frequency analysis (wavelet method using Hanning tapers, width=7, 8-40 Hz with 0.25 Hz resolution, 1 Hz smoothing) followed by spectral parameterization into periodic and aperiodic components (fooof algorithm; Donoghue et al. 2020). The maximum number of peaks and the minimum peak height were set to 4 and 0.1, respectively, to attenuate detection of spurious peaks. Mean R^2^ values obtained during spectral parameterization were 0.90 ± 0.17. Aperiodic-corrected spectra at each time point were averaged in the mu (8-12 Hz) and beta (13-35 Hz) ranges to generate mu and beta power timeseries signals. Scalp channels showing the strongest movement-related reactivity were identified and used for all subsequent analysis (one channel per participant and frequency). This approach allowed separate channels to be identified for mu and beta oscillations. See Figure 2 for depiction of identified channels for each frequency.

**Figure 2.**
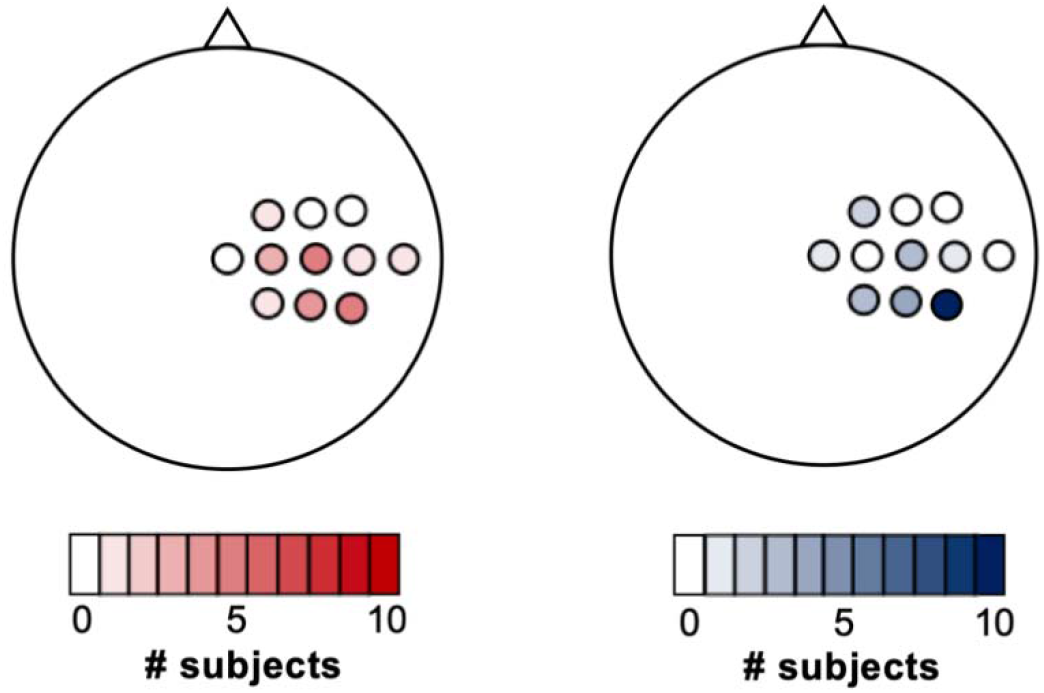
Channels used for analysis of mu and beta oscillatory activity. For mu (left panel), FC1, C6, T8, and CP2 were selected in one subject, C2 was selected in three subjects, CP4 was selected in four subjects, and CP6 and C4 were selected in five subjects. For beta (right panel), Cz and C6 were selected in one subject, FC2 was selected in two subjects, C4 and CP2 were selected in three subjects, CP6 was selected in four subjects, and CP6 was selected in seven subjects.

#### Estimating time of motor command release from M1 (MCR_M1_)

The time of MCR_M1_ was estimated for each finger movement by subtracting the individual MEP latency from each movement’s EMG-defined onset time (see *EMG processing* and Figure 1c-d). Because the MEP latency reflects the amount of time needed for an action potential to travel from M1 to the L. FDI, this approach provided a physiologically-informed estimate of MCR_M1_ required for each finger movement.

#### Phase angle calculation

Pre-processed task-related EEG data were used to identify finger movements containing periodic mu and beta oscillatory activity. Movement-specific aperiodic-corrected time-frequency representations were computed using the same approach described above (see *EEG processing*). For each movement, the power spectra centered on the MCR_M1_ time point (see *Identification of MCR*_*M1*_ *time points*) was used to determine that movement’s dominant mu and beta center frequency.

Movements in which no periodic component with a center frequency in the mu or beta range were present were excluded from further analysis. Data from all remaining movements were band-pass filtered into individually-defined, movement-specific frequency ranges (± 2 Hz relative to each movement’s periodic component center frequency [rounded to the nearest integer]). Movements with EEG activity containing visible artifacts were excluded from further analysis. The phase angle during MCR_M1_ time point was calculated for each movement using the Hilbert transform. Phase angles were combined across all analyzed movements and each subject’s mean phase angle was computed per frequency. As a secondary analysis, phase angles were pooled across all subjects to create a single, group-level phase angle distribution for each frequency band (see *Statistical Analysis*). Overall, 31.01 ± 13.70% of all finger movements contained periodic mu activity at an average of 10.84 ± 0.88 Hz, while 64.94 ± 19.33% of movements contained periodic beta activity at an average of 20.72 ± 4.11 Hz. See Figure 3 for EEG and EMG data and EEG power spectra obtained from a representative subject during the finger movement task.

**Figure 3.**
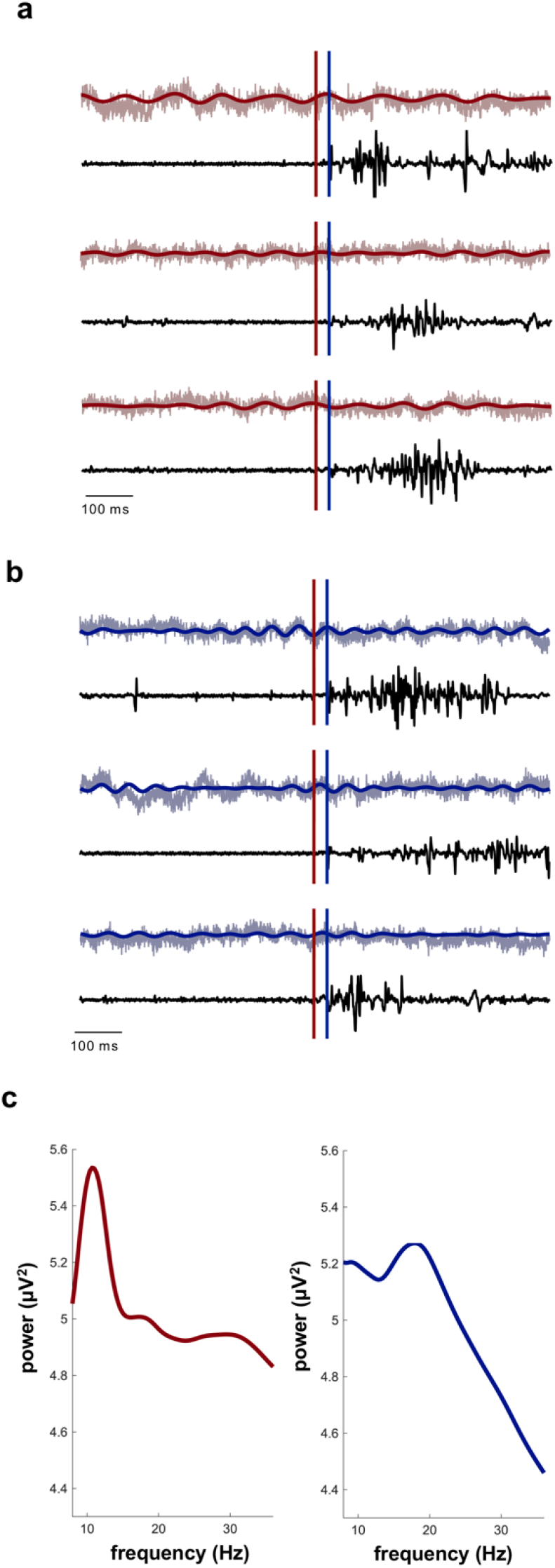
EEG and EMG data from a representative subject. a and b) EEG and EMG data in the mu (a) and beta (b) ranges during three finger movements. Light red (a) and light blue (b) traces indicate raw EEG data, dark red (a) and dark blue (b) traces indicate band-pass filtered EEG data, and black traces indicate raw EMG data. Blue vertical lines indicate EMG onsets and red vertical lines indicate MCR_M1_. Note the oscillatory activity present in all raw EEG traces. c) Aperiodic-adjusted power spectra averaged across all finger movements indicating the presence of mu (left panel, red) and beta (right panel, blue) oscillatory activity.

### Statistical analysis

We tested if phase angles during MCR_M1_ occurred during restricted phases of the mu and beta rhythm by combining the mean phase angles obtained for each subject and frequency during MCR_M1_ across all subjects. Subjects with fewer than 10 trials available for subject-specific mean phase angle calculation (see *Phase angle calculation*) were excluded from statistical analysis (mu, 3 subjects; beta, 1 subject). As a secondary analysis, we pooled phase angles during MCR_M1_ across all subjects within the mu and beta ranges to create a single, large phase angle distribution for each frequency. Phase distributions were tested for unimodal deviations from uniformity using the Rayleigh test. All statistical analysis was performed in MATLAB using custom-written scripts combined with the CircStat Toolbox (Berens et al. 2009) and alpha was equal to 0.05 for all analyses.

## Results

We determined if MCR_M1_ required to produce self-paced voluntary finger movements preferentially occurred during restricted phase ranges of sensorimotor rhythms. Beta phase angles during MCR_M1_ occurred near the peak of the beta cycle (see Figure 4b), exhibiting a significant, unimodal deviation from uniformity at 119.35 ± 144.92° (p=0.002). Similarly, the pooled group-level distribution of beta phase angles during MCR_M1_ also exhibited a significant unimodal deviation from uniformity at 123.94 ± 168.98° (p=0.008). In contrast, phase angles during MCR_M1_ were uniformly distributed across all angles of the mu cycle, with neither the mean group-level distribution or the pooled group-level distribution of mu phases during MCR_M1_ significantly deviating from uniformity (see Figure 4a, p>0.25 for both).

**Figure 4.**
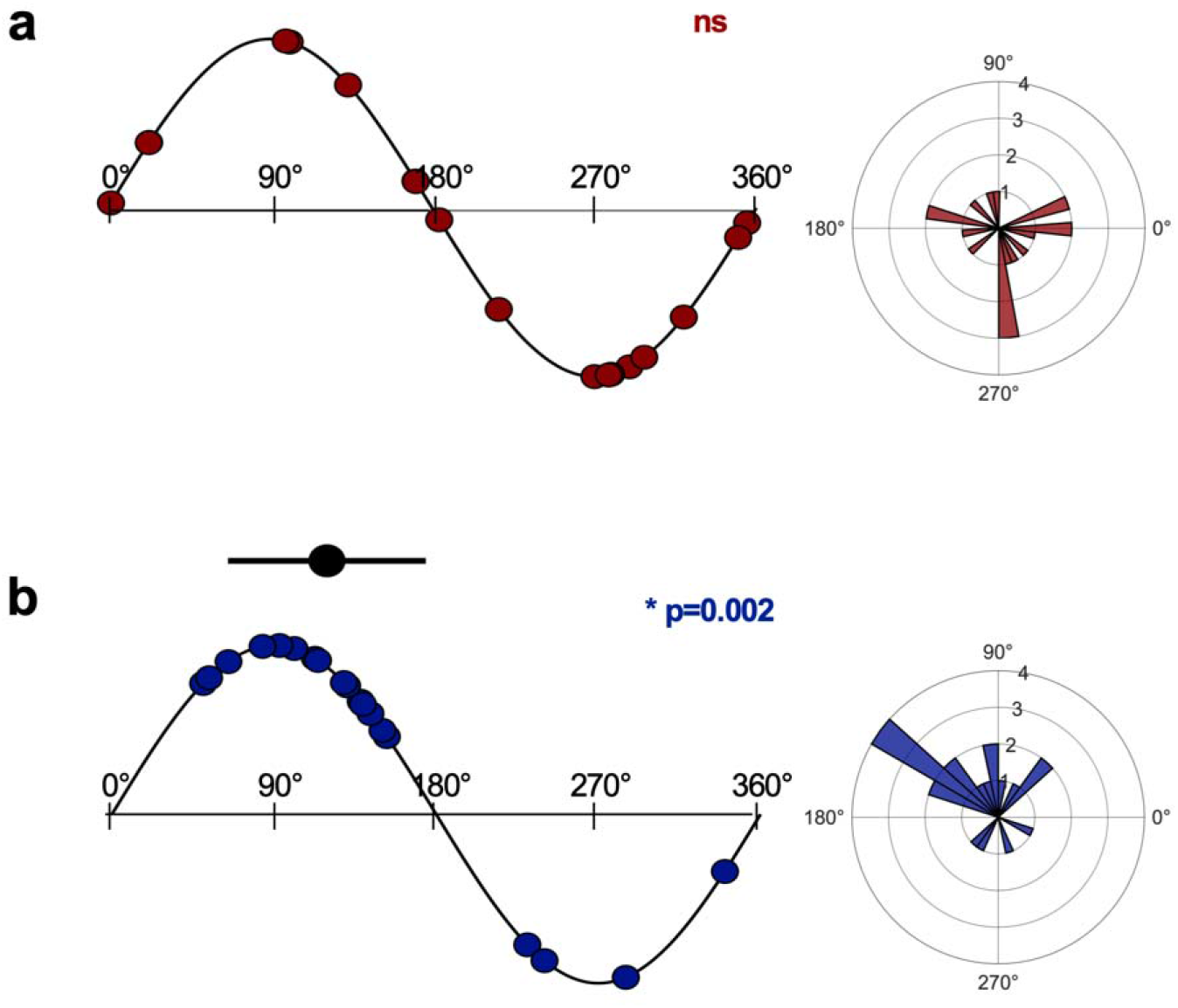
Mu and beta phase angles during MCR_M1>_. Mu (a) and beta (b) phase angles during MCR_M1_ along the oscillatory cycle (left panel) and in phase space (right panel). For left panels, each dot indicates a single subject’s mean phase angle during MCR_M1_. Right panels contain phase angle histograms representing group-level distributions of mean phase angles during MCR_M1_. Note that phase angles during MCR_M1_ were evenly distributed across the mu cycle (a) but occurred near the peak of the beta cycle at 119.35° (b). Radius values indicate the number of subjects showing mean phase angles at a given phase bin. The black dot and horizontal line reflect the mean beta phase angle and the group-level standard deviation. Mean and standard deviations are not shown for mu due to lack of significant deviations from uniformity. *reflects significance at p<0.05. ns reflects p>0.05.

## Discussion

In this study, we evaluated whether motor commands required to produce self-paced finger movements were preferentially released from M1 during restricted phase ranges of sensorimotor mu and beta rhythms. We report that phase angles during MCR_M1_ were preferentially clustered at ~120° of the beta cycle but were evenly distributed across the mu cycle.

Our finding that MCR_M1_ occurred during restricted phase ranges of the ongoing beta cycle is consistent with previous work reporting that sensorimotor beta oscillatory activity covaries with M1 single-neuron spiking rates (Murthy and Fetz 1996a; Zanos et al. 2018), M1 population-level neuronal activity (Miller et al. 2012), and corticospinal output (Khademi et al. 2018; Keil et al. 2013; Torrecillos et al. 2020), all of which are necessary for voluntary movement. In addition, it has been proposed that corticospinal communication during movement occurs through phase-synchronization between cortical and spinal oscillatory activity (i.e., corticomuscular coherence; Farmer et al. 1993; Conway et al. 1995; Mima and Hallett 1999; Womelsdorf et al. 2007). Outside of the motor domain, perceptual function is enhanced at specific phases along oscillatory cycles recorded from task-relevant brain regions (Busch et al. 2009; Dugué et al. 2009; Hanslmayr et al. 2013; Baumgarten et al. 2015; Busch and VanRullen 2010; VanRullen 2016; but see also Ruzzoli et al. 2019), suggesting that human perception occurs through rhythmic sampling of the environment (VanRullen 2016). Our results show for the first time that the release of motor commands from M1, a process essential to voluntary movement, exhibits similar rhythmicity.

Motor commands were preferentially released from M1 at ~120° of the ongoing beta cycle but were released uniformly across the mu cycle. Why might motor command release be coupled to the beta but not mu rhythm, and why around 120°? Mu and beta rhythms are generated by distinct neural mechanisms (Stolk et al. 2019). Mu activity localizes to the primary somatosensory cortex (Salmelin and Hari 1994), exhibits little if any somatotopy (Salmelin et al. 1995) and travels caudo-rostrally across the cortex (Stolk et al. 2019), while beta activity localizes to the primary motor cortex (Salmelin and Hari 1994), exhibits more precise somatotopic organization (Salmelin and Hari 1994; Salmelin et al. 1995) and travels rostro-causally across the cortex (Stolk et al. 2019). Beta activity is closely tied to movement initiation, as entraining beta oscillations slows movement (Pogosyan et al. 2009) and patients with Parkinson’s disease often exhibit exaggerated beta activity that correlates with bradykinesia and movement initiation deficits (Little and Brown 2014; Martin et al. 2018). Further, corticomuscular coherence is strongest in the beta range (Conway et al. 1995; Baker et al. 1997; Mima and Hallett 1999), indicating the presence of a beta-specific communication channel between M1 and spinal motoneurons (Romei et al. 2016; van Elsjiwk et al. 2010; Khademi et al. 2018) with EMG bursts locked to trough phases of sensorimotor beta activity (Baker et al. 1997; Mima and Hallett 1999). For motor commands to produce EMG bursts that coincide with trough phases, they would need to be released between 90-125° (assuming a 20 Hz rhythm oscillating at ~7.2° per ms, as seen here, see *Phase angle calculation*) which is consistent with the ~120° beta angle identified here. This phase range also coincides with the peak of the beta cycle, during which M1 single-neuron spiking rates and population-level neuronal activity are at their lowest (Murthy and Fetz 1996a; Miller et al. 2012). When combined with this previous work, our results suggest that decreased background neuronal activity at the peak of the beta cycle may increase the signal-to-noise ratio of motor commands, allowing them to be efficiently transmitted from M1 to spinal motoneurons.

In conclusion, we report that motor commands were preferentially released from M1 near the peak of the beta cycle at ~120° but were released uniformly across the mu cycle during a self-paced voluntary finger movement task. Results are consistent with the notion that endogenous sensorimotor beta phase actively shapes release of motor commands from the healthy human brain.

## Acknowledgements

Data included in this publication were acquired at the Human Cortical Physiology and Neurorehabilitation Section (directed by Leonardo Cohen) at NINDS.

